# Donor Age and Time in Culture Affect Dermal Fibroblast Contraction in a Hydrogel Skin Graft Model

**DOI:** 10.1101/2021.11.29.469875

**Authors:** Amber Detwiler, Kathryn Polkoff, Lewis Gaffney, Donald Freytes, Jorge Piedrahita

**Affiliations:** Comparative Medicine Institute, North Carolina State University, Raleigh, NC; Joint Department of Biomedical Engineering, North Carolina State/ University of North Carolina-Chapel Hill, Raleigh NC; Department of Molecular Biomedical Sciences, North Carolina State University, Raleigh, NC

**Keywords:** cell proliferation, collagen, dermis, extracellular matrix, fibroblasts, hydrogels, skin, wound healing

## Abstract

Regenerating functional skin without the formation of scar tissue remains an important goal for Tissue Engineering. Current hydrogel-based grafts minimize contraction of full-thickness skin wounds and support skin regeneration using adult or neonatal foreskin dermal fibroblasts, which are often expanded *in vitro* and used after multiple passages. Based on the known effects of 2D tissue culture expansion on cellular proliferation and gene expression, we hypothesized that differences in donor age and time in culture may also influence the functionality of 3D skin constructs by affecting fibroblast-mediated graft contraction. To validate these predicted differences in fibroblast phenotype and resulting 3D graft model contraction, we isolated porcine dermal fibroblasts of varying donor age for use in a 2D proliferation assay and a 3D cell-populated collagen matrix contractility assay. In 2D cell culture, doubling time remained relatively consistent between all age groups from passage 1 to 6. In the contractility assays, fetal and neonatal groups contracted faster and generated more contractile force than the adult group at passage 1. However, after 5 passages in culture, there was no difference in contractility between groups. These results show how cellular responses differ based on donor age and time in culture, which could account for important differences in biomanufacturing of 3D hydrogel-based skin grafts. Future research and therapies using bioengineered skin grafts should consider how results may vary based on donor age and time in culture before seeding.

**IMPACT STATEMENT:** Little is known about the impact of donor age and time in culture on the contraction of the 3D hydrogel-based graft. These results show how cellular phenotypes differ based on donor age and time in culture, which could account for important inconsistencies in biomanufacturing of skin grafts and *in vitro* models. These findings are relevant to research and therapies using bioengineered skin graft models and the results can be used to increase reproducibility and consistency during the production of bioengineered skin constructs. Future *in vivo* studies could help determine the best donor age and time in culture for improved wound healing outcomes or more reproducible *in vitro* testing constructs.

## INTRODUCTION

Due to the vital functional role skin plays in protection, insulation, and prevention of water loss, severe skin injuries such as burns and deep wounds can be debilitating and life-threatening ^1,2^. Skin grafting is performed annually in 30% of burn hospitalizations and as treatment for chronic skin wounds ^3,4^. While autologous skin transplants for full-thickness skin wounds is common, it is not a viable treatment for larger, more severe wounds due to insufficient donor material to cover the affected area. Engineered biomaterials can substitute autologous grafts, thus eliminating the need for harvesting autologous tissues while also supporting more natural healing. Many bioengineered grafts utilize a hydrogel-cell construct, which focuses on replicating the dermis, the innermost layer of skin composed of dermal fibroblasts in a network of collagen and elastic fibers, by isolating and seeding dermal fibroblasts in a collagen-based hydrogel. These hydrogels can be created as a single-layer or bi-layer dermal graft. While there are many acellular skin grafts on the market, they do not address the poor barrier function of grafts without a cellular component, while also promoting ECM deposition, secreting growth factors, and lengthening the time that the graft covers the wound ^5^. Traditionally, patient-derived dermal fibroblasts are isolated, expanded and used for autologous grafts. Another common approach is the use of neonatal foreskin dermal fibroblasts given their availability and perceived notion that they will be more resilient and promote faster healing due to their neonatal phenotype. A summary of skin grafts and skin models currently on the market, including cell age and time in culture at seeding, is shown in Table 1.

Even with differences in donor age (neonate to adult with no definition of adult age) and time in culture (passage 1-8) at seeding among grafts and skin models (Table 1), little is known about how these two variables impact the contractile properties of the 3D graft, and thus their eventual utility for wound healing applications. We hypothesized that dermal fibroblasts of differing donor age and time in culture result in different graft properties as predicted by results using collagen hydrogels.

This hypothesis was based on differences in levels of regeneration, scarring, and contraction observed *in vivo* during healing between fetal, neonatal, and adult skin, as well as the impact of increased 2D culture on cellular morphology and phenotype. *In vivo* wound healing studies show differences between adult and neonatal skin, with neonates exhibiting faster re-epithelialization, reduced inflammation and scarring, and better regeneration of natural tissue structure ^19^. *In vivo* studies have also shown that fetal skin can undergo scarless-healing, contrary to comparably wounded adult skin which cannot heal to its original state and instead forms scar tissue lacking skin appendages ^20^. Two-dimensional culture does not accurately mimic conditions *in vivo* and increased time in culture in this environment has been found to disturb cell-matrix interactions and even alter cell morphology, division method, and cell polarity ^21^. Because of all these properties, we hypothesized that donor age and increased time in culture may affect cellular response when seeded in a hydrogel-based skin graft model. A porcine model was chosen based on research by Sullivan et al. which found that porcine wound healing models agree with human studies 78 percent of the time, whereas small-animal studies agree only 53 percent of the time ^22^. Porcine skin is similar to human skin in thickness, hair distribution, and coat attachment ^22^.

In order to examine the phenotypic behavior of dermal fibroblasts in collagen-based hydrogels, changes in contractile and growth properties of skin graft models containing fibroblasts of varying donor age and time in culture were measured. Understanding the impact of these factors on skin grafts can improve wound healing outcomes and increase reproducibility across skin grafts and models, leading to more consistent, reliable results.

## MATERIALS AND METHODS

### Primary cell isolation

Porcine dermal fibroblasts were isolated from fetal (isolated at day 42 of pregnancy), neonatal (less than 24 hours after birth), and adult (over 6 months) dermis in this study. This study was carried out in strict accordance with the recommendations in the Guide for the Care and Use of Laboratory Animals of the National Institutes of Health ^23^ and approved by the Institutional Animal Care and Use Committee of North Carolina State University.

Dermis was removed from gestational day 42 fetal pigs and the tissue was then digested in a 0.025% trypsin and 0.5% ethylenediaminetetraacetic acid solution in a 37 degree Celsius (°C) tumbling incubator for 45 minutes. An equal volume of alpha Dulbecco’s Modified Eagle Medium (DMEM) medium was added to resuspend the cell suspension in medium. Cells were cultured at 37 °C for at least 3-5 days, checking for attachment and adding media if necessary, and then split after 6-10 days. At 80 percent confluence, the fetal fibroblasts were harvested and frozen in dermal freezing media (6:3:1 ratio of DMEM, fetal bovine serum (FBS), and dimethyl sulfoxide respectively).

Adult and neonatal dermal fibroblasts were isolated by first removing fat and shaving hair from a fresh porcine skin sample and dicing prepared skin into 2-3 millimeter square pieces and incubated at 4 °C overnight in a 10 mg dispase per milliliter (mL) of 1X Dulbecco’s Phosphate Buffered Saline (DPBS) solution. After manually separating the dermis from the epidermis, the dermis was digested in 4 mg collagenase per 1 mL DMEM in a rolling incubator at 37 °C for 1-3 hours. After vortexing the dermis in solution in three, 10 second intervals, the liquid was filtered with a 100 micrometer (μm) filter and frozen in dermal freezing media and stored at −80 °C before transfer to liquid nitrogen until use.

### Proliferation assay

For the proliferation assay, cells were seeded at a known cell density of 2.2×10^5^ in a 6-well plate. Fibroblasts were passaged with 0.05% trypsin when confluent and counted with an automatic cell counter. Doubling time was then calculated using the following formula:

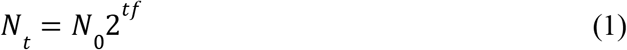

where *N_t_* is the number of cells at passage, *N*_0_ is the number of cells seeded initially, *t* is the time in hours from seeding to passage, and 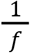 is the doubling time in hours. Doubling times were calculated using cells that had undergone at least one passage after thawing (apart from passage 1). This assay was performed with at least two biological replicates and three technical replicates at each age and passage.

### Hydrogel creation

For every 1 mL of a hydrogel of ECM concentration 6 mg/mL the following were combined on ice: 600 μL Bovine Type-1 Collagen BioMatrix, 275 μL 1X PBS, 67 μL 10X PBS, and 5 μL phenol red as a pH indicator. This mixture was vortexed to combine before adding 90 μL of 0.1M sodium hydroxide and vortexing again, changing the color from the yellow, acidic pH to a pink, neutral pH. To seed the hydrogel with cells, the gel was diluted to an ECM concentration of 3 mg/mL by adding an equal volume of cell suspension in DMEM. This process is simplified in Figure 2A. The gels used in this study were seeded at a density of 1.5×10^6^ cells per mL. This mixture was vortexed to combine and plated accordingly.

### Contraction assay

For the contraction assay, the hydrogel/cell mix described above was plated 0.5 mL per well in a 24-well plate, avoiding bubbles. The hydrogels were incubated at 37 ° C for 15-30 minutes, or until set, before gently detaching the gel from the well sides with a pipette tip and adding 1 mL DMEM with 15% FBS and 1% antibiotic/ antimycotic. The contraction assay was repeated with fibroblasts isolated from at least two different pigs of the same age and cultured to the same passage (either passage 1 or 5) before seeding. The assay for each sample had at least two technical replicates. One well was seeded without any cells and only collagen hydrogel in order to act as a control, with no contraction expected in this well.

Photos were taken every day for 5-7 days after seeding. Percent contraction as determined by surface area was calculated in ImageJ and MATLAB was used to calculate the day at which each sample reached a threshold of 70 percent of total contraction.

### Micropost assay

A micropost assay was used to measure force generation of the cells in the hydrogel, as done by Nandi et al ^24^. Microposts were created using polydimethylsiloxane in a 3D printed mold. These microposts were then plasma treated and stored in deionized water. Right before use, the microposts were sterilized with 70% ethanol and set out to air dry under a hood. The microposts were then coated with 0.3% bovine serum albumin for 10 minutes, aspirated to dry, and seeded with hydrogel. Approximately 1.5 μL of hydrogel of the same cell density and ECM concentration of the contraction assay was pipetted into each micropost. The gels were incubated for 15-30 minutes, or until set, before gently adding 1 mL DMEM with 15% FBS and 1% antibiotic/ antimycotic to the microposts in the well of a 24-well plate. Since the same hydrogel/cell mix was used in the micropost assay as was used in the contraction assay, the hydrogel-only well described in the contraction assay protocol also acted as a control for the micropost assay. The micropost assay was performed with at least two biological replicates and at least two technical replicates.

Photos were taken every day for 5-7 days after seeding. Force generation was calculated with the equation F=kδ, where F is the force in micronewtons (μN) applied to cause the micropost deflection, k is the spring constant of the microposts, and δ is the deflection distance (μm). ImageJ was used to measure deflection, and the constant k was assumed to equal 7.5 newtons per meter ^24^.

### Gene expression analysis

RNA extraction was performed using a Zymo RNeasy Mini-Prep Kit. Dermal fibroblasts were suspended in lysis buffer immediately after passages 1 and 5 according to the manufacturer’s instructions with the on-column DNAse digest. cDNA was synthesized according to the Agilent AffinityScript cDNA Synthesis Kit protocol. PCR products were run using gel electrophoresis to confirm that product sizes matched those shown in Supplemental Table I. For gene expression analysis, each of the genes in Supplemental Table I was measured by RT-qPCR in the qTOWER3 by Analytik Jena with the iTaq Universal SYBR Green Supermix kit with GAPDH as the control. ddCt values were then normalized to passage 1 for each age group (i.e. adult passage 5 was normalized to adult passage 1, neonatal passage 5 to neonatal passage 1, etc.). Three biological replicates were analyzed for each age and passage number.

### Statistical analysis

Micropost assay and contraction assay data were analyzed using an unpaired, two-tailed t-test. The qPCR data were analyzed using Welch’s t-test for comparison between passage 1 and passage 5 data within each age group in GraphPad Prism. Significance was set at P<0.05.

## RESULTS

### Measured proliferation was consistent across cell ages

Dermal fibroblasts isolated as shown in Figure 1A were frozen immediately after isolation at passage zero before being thawed and plated for use in the proliferation assay. No morphological differences in fibroblasts based on age were noted (Figure 1B-D). All age groups decreased in doubling time with a smaller standard deviation immediately after passage 1 and into passage 2 (Figure 1E). No other notable differences in doubling time were noted between age groups and passages, indicating consistent proliferation from passages 1 to 6 as measured by doubling time.

**Figure 1.**
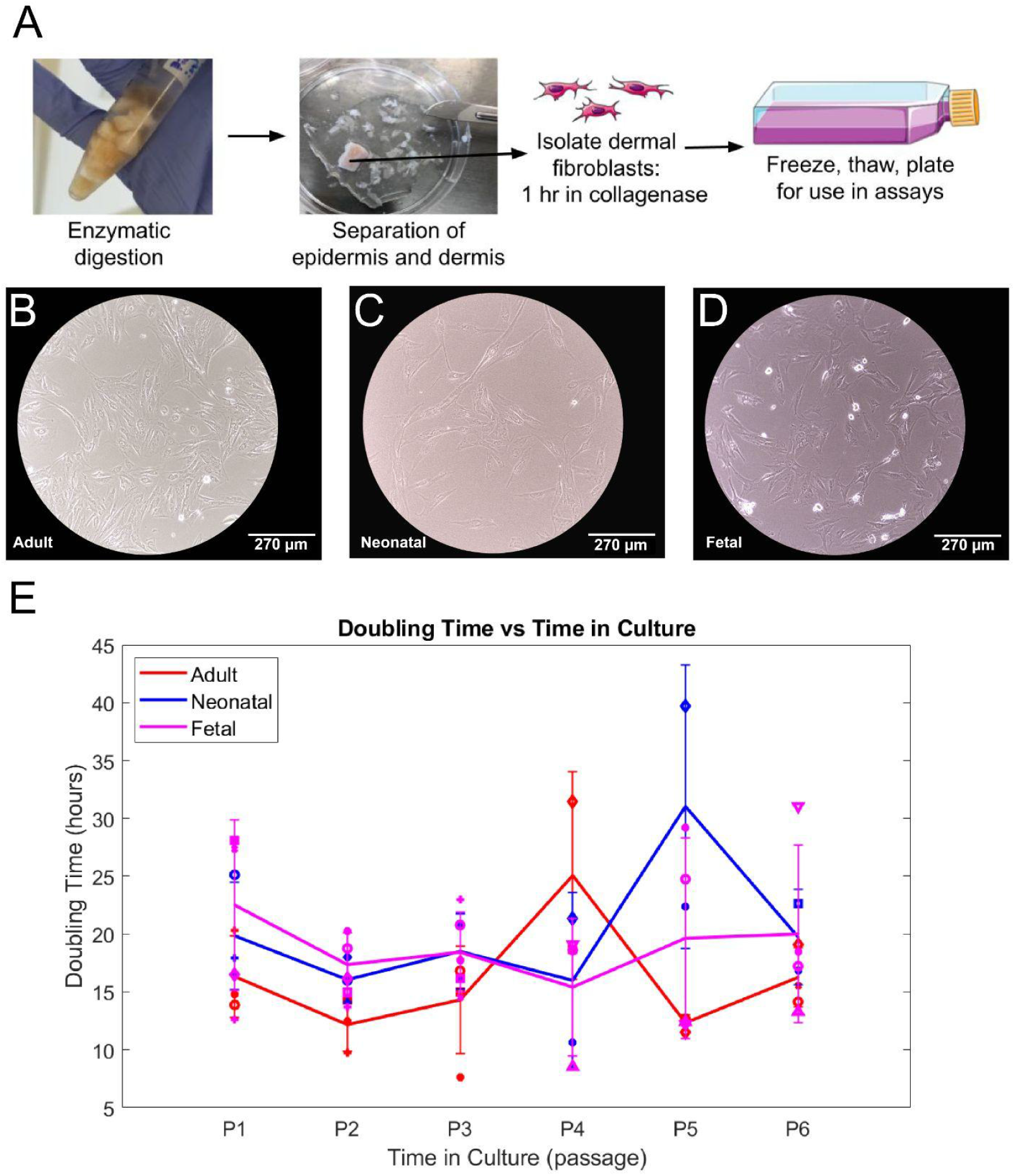
Proliferation measured by doubling time remained relatively consistent between adult, neonatal, and fetal fibroblasts at passages 1-6. (A) Schematic illustration of primary cell isolation of dermal fibroblasts and epidermis from porcine skin. (B-D) Brightfield 10x image of passage 1 adult, neonatal, and fetal dermal fibroblasts in 2D culture. (E) Average doubling time in hours for adult (red), neonatal (blue), and fetal (magenta) samples from passage 1 to passage 6. Each marker shape corresponds to a different sample for that age group. Cells from at least two different pigs were analyzed at each passage, with three technical replicates of each cell sample averaged to produce each point above.

### Adult fibroblasts contract collagen gel less than other ages at passage 1

To determine the differences in contractile properties across age groups after minimal culture, we first measured the impact of donor age upon the rate of contraction of collagen hydrogels. After seeding the dermal fibroblasts within a collagen hydrogel as shown in Figure 2A, the change in area was measured over the course of five days. The adult dermal fibroblasts (Figure 2C) took significantly longer time than the other two groups to reach the contractile threshold of 70 percent of total contraction (Figure 2D-E).

**Figure 2.**
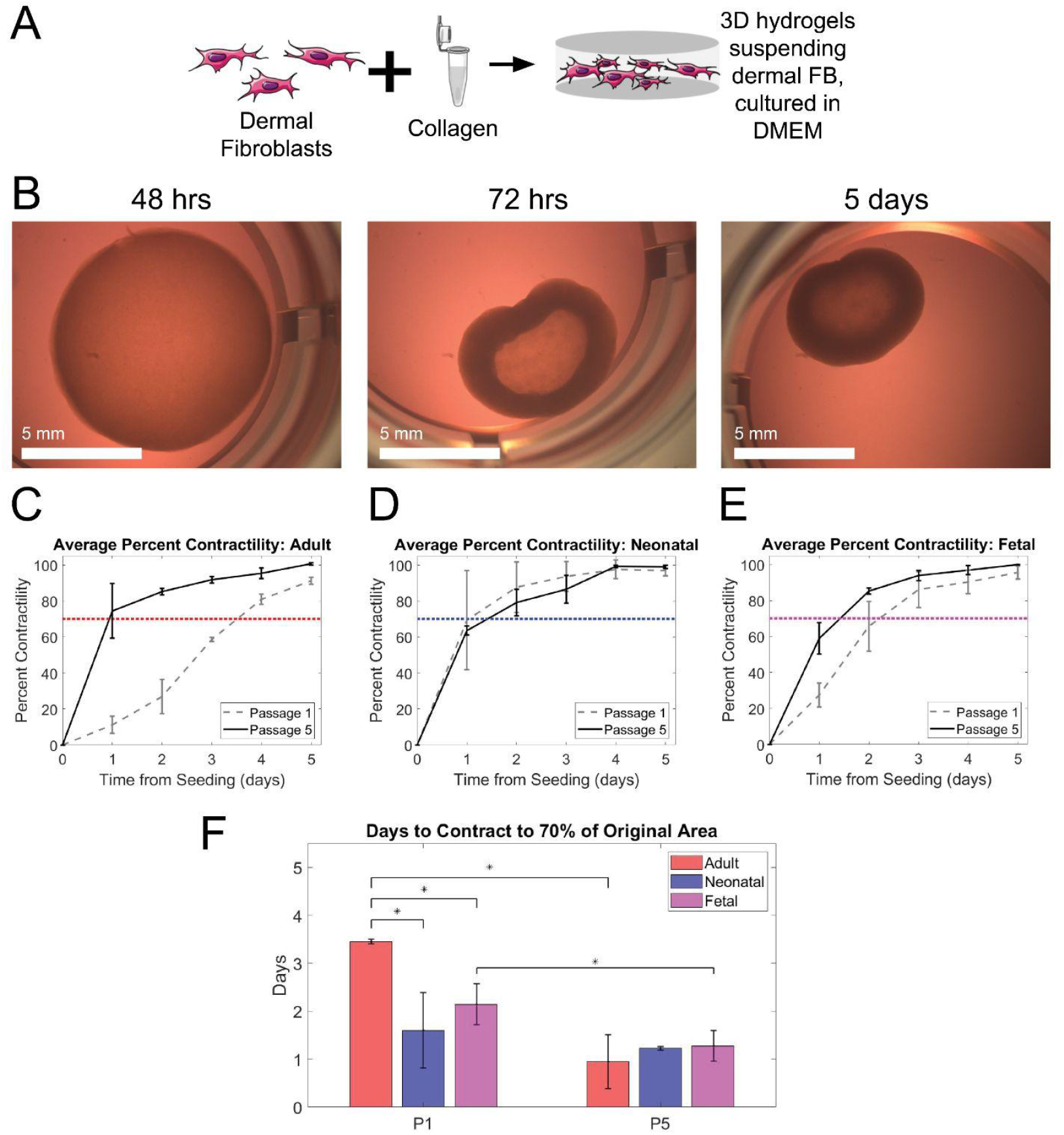
Adult dermal fibroblasts contracted slower than neonatal and fetal when seeded at P1. Adult, neonatal, and fetal age groups experienced similar contraction between age groups, and faster contraction within age groups when seeded at P5. (A) Schematic illustration displaying the combining of dermal fibroblasts and bovine type I collagen gel to culture in 3D. (B) Fetal fibroblasts seeded in collagen contractility assay at 48 hours, 72 hours, and 5 days after seeding. (C-E) Time in days for each age group to contract to a threshold of 70 percent, grouped by the passage number of cells at seeding (passage 1 on left and passage 5 on right). Statistical significance between groups with an alpha value of 0.05 is indicated by an * and bars indicate standard deviation. (F) Average percent contractility from Day 0 (seeding) to Day 5 normalized to maximum contraction. Adult (left, red), neonatal (middle, blue), and fetal (right, magenta) shown. Data for cells seeded at passage one are shown in dashed grey and those seeded at passage five are shown in black. A threshold of 70 percent is shown in dotted red, blue, and magenta for adult, neonatal, and fetal respectively. Cells from at least two different pigs with three technical replicates were analyzed in each group.

To quantitatively confirm the effectiveness of the contraction assay in accurately measuring contractile forces, force generation was measured using a micropost assay (Supplemental Figure 1A). Neonatal and fetal dermal fibroblasts generated more force than adult fibroblasts, although only fetal and adult were found to be statistically significantly different (Supplemental Figure 1B). Based on the similar contractile force generation results of the contraction assay and micropost assay at passage 1, the contraction assay alone was deemed sufficient to measure contractile forces for samples seeded at passage 5.

### Contraction was similar between age groups and faster within groups at passage 5

Since many of the products on the market expand the fibroblasts in culture (Table I) before using, the next step was to determine how serial passaging of the fibroblasts would impact hydrogel contraction. Passage 5 was chosen as the average of the passages provided in Table I. Results from contraction of passage 5 cells overlaid with passage 1 cells show that the neonatal and fetal donors have similar contraction curves at both passages, but the adult dermal fibroblasts contracted faster at passage 5 than passage 1 (Figure 2C).

**Table I.** Examples of hydrogel-based skin grafts and skin models on the market. Dermal fibroblast cell type (allogenic or autologous), cell age (adult and/or neonatal), time in culture at seeding (passage number) are shown for comparison.

| Graft | Manufacturer | Cell Type | Cell Age | Time in Culture at seeding | Singular-layer/bi-layer | Biomaterial | Source(s) |
| --- | --- | --- | --- | --- | --- | --- | --- |
| *Transcyte* | Advanced Tissue Sciences | Allogenic | Neonatal | Passage 8 | Singular-layer | Nylon mesh with a silastic layer | <sup>6</sup> |
| *Apligraf* | Organogenesis Inc | Allogenic | Neonatal | Passage 5-7 | Bi-layer | Bovine collagen type I | <sup>7</sup> |
| *Dermagraft* | Organogenesis Inc | Allogenic | Neonatal | Passage 8 | Singular-layer | Polyglycolic acid mesh | <sup>8</sup> |
| *MyDerm* | Universiti Kebangsaan Malaysia | Autologous | Adult | Passage 3 | Bi-layer | Plasma-fibrin | <sup>9,10</sup> |
| *OrCel* | Forticell Bioscience, Inc. | Allogeneic | Neonatal | Passage 4 | Bi-layer | Type I bovine collagen sponge | <sup>11,12</sup> |
| *PermaDerm* | Regenicin | Autologous | Adult | Passage 2 | Bi-layer | Absorbable collagen substrate | <sup>13</sup> |
| *StrataGraft* | Stratatech Corp. | Allogenic | Adult or Neonatal | Not specified | Bi-layer | Murine collagen | <sup>14,15</sup> |
| Model | Manufacturer | Cell Type | Cell Age | Time in Culture at seeding | Singular-layer/bi-layer | Biomaterial | Source(s) |
| *Phenion* | Henkel | Allogenic | Neonatal | Passage 1 | Bi-layer | Not specified | <sup>16,17</sup> |
| *EpiDerm-FT* | MatTek Corp. | Allogenic | Adult or Neonatal | Not specified | Bi-layer | Not specified | <sup>18</sup> |

Comparison of contraction rate (time required for gel to contract to 70% of total contraction) showed that fetal contracted faster than adult (p=0.001) and neonatal faster than adult (p=0.02) (Figure 2F). When seeded at passage 5, however, there were no statistically significant differences between any age group. This is because, while the neonatal group contraction did not change between P1 and P5 (p=0.57), the adult and fetal groups demonstrated significantly increased contraction after 5 passages (p=0.004 and 0.04, respectively). These results suggest that although there were significant differences in contraction of early passage cells based on donor age, after 5 passages their behavior was more homogenous and suggests that time in culture generally increases contractile properties of fibroblasts.

### Effect of passage on *TAZ, STAT1*, and *COL1A2* expression

To further understand differences in the cellular phenotype that could contribute to the contraction observed in Figure 2, gene expression analysis was performed focusing on 9 genes involved in wound healing, contraction, and ECM deposition.

From passage 1 to passage 5, TAZ fold change showed a statistically significant difference within adult, neonatal, and fetal groups with p-values of 0.002, 0.03, and 0.02 respectively (Figure 3A). STAT1 significantly increased from neonatal passage 1 to neonatal passage 5 with a p-value of 0.03 (Figure 3B). The adult group saw a significant increase from passage 1 to 5 for fold change in expression of COL1A2 with a p-value of 0.02 (Figure 3C).

**Figure 3.**
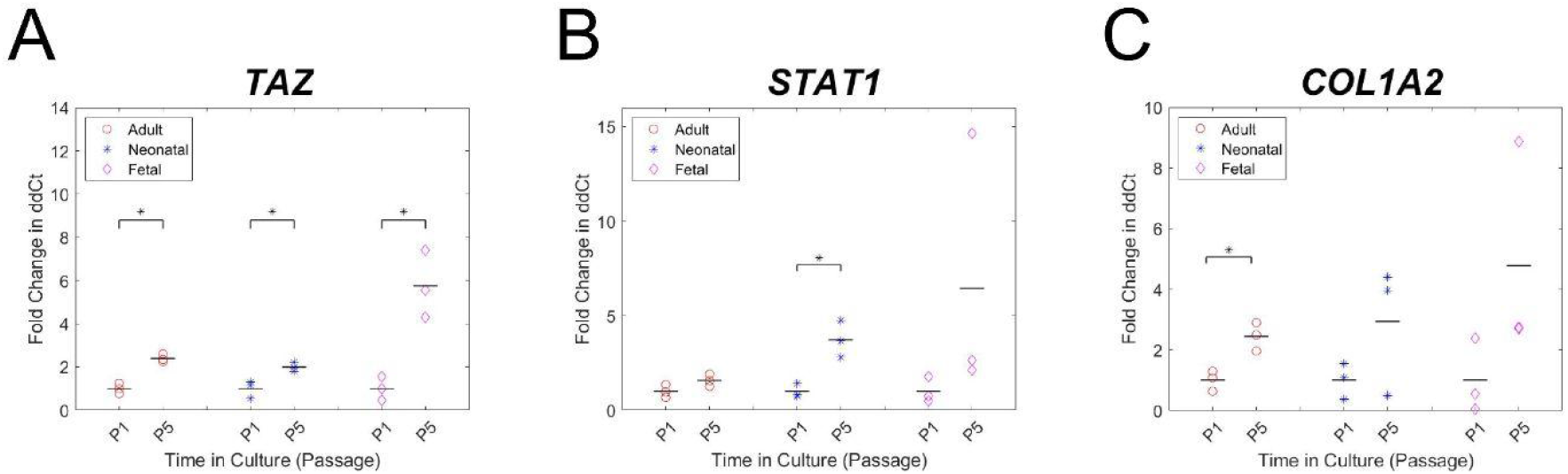
Gene expression analysis for a panel of genes related to wound healing, contraction, and ECM deposition. (A-C) Fold change in ddCt from passage 1 to 5 for adult (red circle), neonatal (blue cross), and fetal (magenta diamond) dermal fibroblasts. The mean for each group is indicated by a black bar. dCt values were standardized to GAPDH, followed by ddCt values where adult passage 5 was standardized to adult at passage 1, neonatal passage 5 to neonatal passage 1, etc. Cells from at least two different pigs with three technical replicates were analyzed in each group. Statistical significance between groups with an alpha value of 0.05 is indicated by a black bar and *

**Figure 4.**
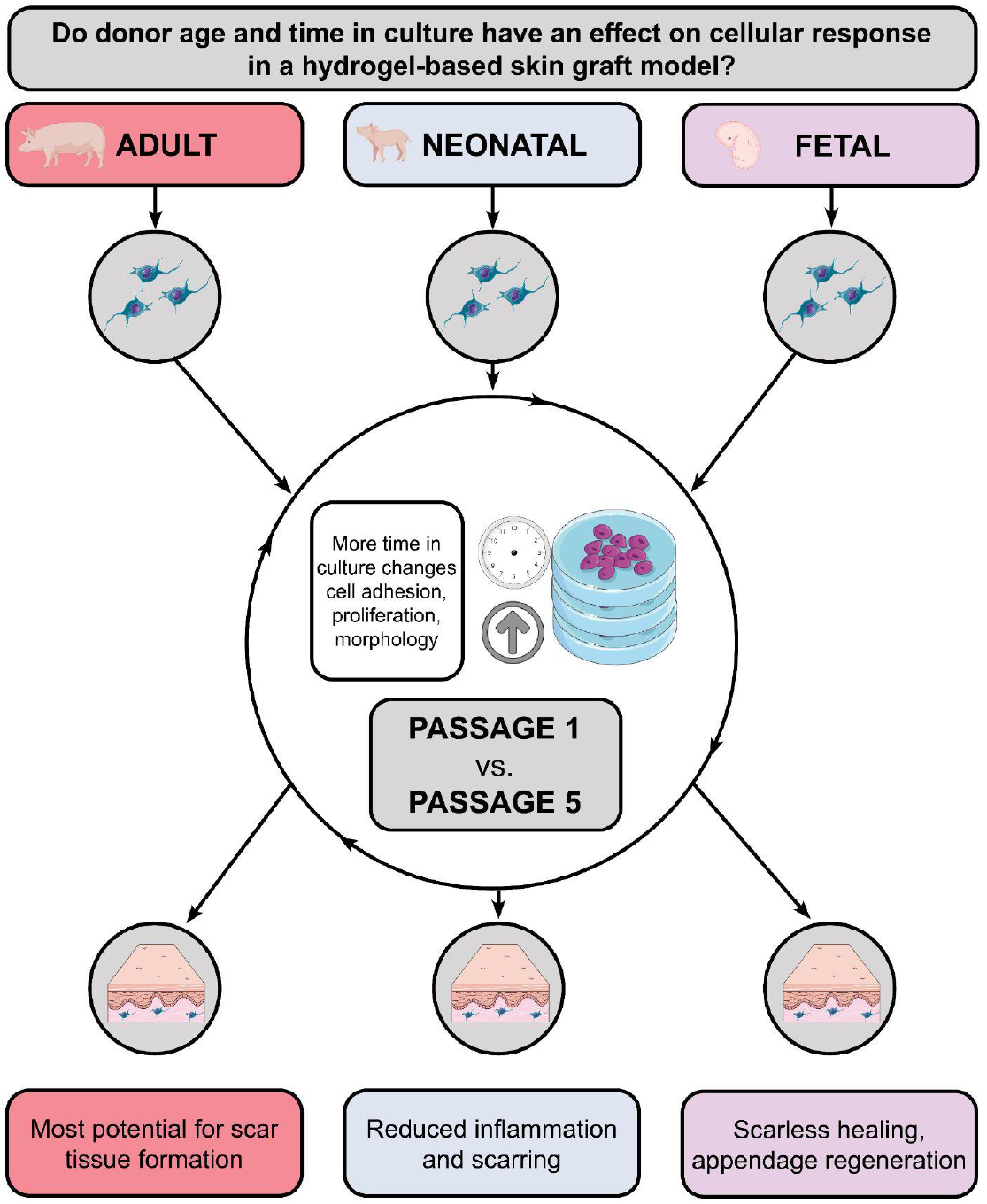
Graphical abstract of research question and hypotheses. Donor ages of isolated dermal fibroblasts are divided into adult (red), neonatal (blue), and fetal (magenta) groups before culturing for varied amount of time (until passage 1 or passage 5). Phenotypic outcomes were measured in a hydrogel-based skin graft model. Hypotheses regarding resulting wound healing outcomes are matched by color at the bottom.

## DISCUSSION

The focus of this research was to understand how donor age and time in culture, which vary in existing skin grafts and models, affect fibroblast behavior and contraction of a fibroblast-populated collagen matrix. Understanding these differences could be beneficial in manufacturing a more consistent and predictable skin graft model.

Doubling time remained relatively consistent between all age groups across the first 6 passages, suggesting that increased expansion on tissue culture plastic did not have any major effects on fibroblast proliferation rates. Based on this fibroblast proliferation and growth rate, there appears to be no advantage to expanding one donor age or passage over another. However, gene expression analysis of donor cells found *TAZ, STAT1*, and *COL1A2* expression to significantly increase from passage 1 to passage 5 in some groups, suggesting that increased time in culture before seeding may affect graft/skin model behavior given the potential phenotypic changes suggested by the measured gene expression. This phenotypic change should be explored in more detail in future studies.

Contraction was similar between age groups at passage 5 but not at passage 1. These results are supported by Hopp et al. which found substrate stiffness to affect adhesion, proliferation and morphology, meaning that increased time in culture on stiff tissue culture plastic causes the cells to behave differently when moved to the much less stiff hydrogel environment ^25^. Whereas the time to reach 70% contraction decreased significantly from passage 1 to passage 5 in the adult model, there was no significant difference in the neonatal model from passage 1 to passage 5. This could suggest that neonatal donor cells present more a consistent phenotype and predictable contraction at a range of passages. Alternatively, adult donor fibroblasts may allow for tunable contraction based on time in culture before seeding.

From a manufacturing standpoint, these results could also help explain some of the variation of fibroblast properties and therefore use this knowledge to engineer a more consistent skin graft product. The similar contraction behavior among age groups at passage 5 as opposed to passage 1 may offer more consistency in skin graft or skin model outcomes, more usable cells per donor tissue, and more predictable scaffold needs. Overall, these results provide insight into fibroblast behavior in a skin graft model based on age and time in culture, factors which may also apply to graft outcomes when applied *in vivo*.

Current skin grafts are known to reduce wound contraction and scarring compared to healing by secondary intention; however, contraction after initial skin graft application can create a need for surgical release and application of additional skin graft ^26^. For skin wounds near joints, this can increase the risk for graft failure and total cost of treatment ^26^. Similarly, *in vitro* skin model contraction can result in detachment of the epidermal layer from the insert wall, creating a less-functional skin model ^27^. Research has focused on graft contraction based on matrix formation and properties, including the effects of temperature on plasma-based hydrogel contraction ^28^, and reduced graft contraction with cross-linked collagen ^27^ and nanofibrillar cellulose ^29^ hydrogels.

While a fibroblast-populated collagen matrix is by no means a full measure of the complete nuances of *in vivo* wound healing contraction, the different contractile properties based on donor age and time in culture could foreshadow different graft outcomes based on the characteristics of the cells seeded in the graft. Cellular phenotype as measured by proliferation, protein synthesis, contraction, and cell viability has been shown to be similar between *in vivo* wound healing models and the *in vitro* fibroblast-populated collagen matrix model ^30^. The contraction of the fibroblast-populated collagen matrix is an indicator of cell activation, with this behavior associated with *in vivo* wound healing ^31^. This research not only shows differing cellular phenotypes *in vitro* based on donor age and time in culture, but could suggest different *in vivo* wound healing outcomes due to these varying cellular phenotypes.

Future assays expanding on these results should assess gene expression at a broader range of cell donor ages and time in culture and compare the cell types discussed *in vivo*. This research is useful for all current and future applications of hydrogel-based skin grafts and could also aid in determining which donor age is best for supporting skin appendage regeneration in biomaterial skin grafts.

## CONCLUSIONS

Findings show that in 2D cell culture, doubling time remained relatively consistent between all age groups from passage 1 to 6. Contraction of collagen hydrogels differed significantly based on donor age as well as time in culture at seeding. Gene expression analysis of donor cells showed changes in *TAZ, STAT1*, and *COL1A3* with increased time in culture. These results provide support for differing cellular phenotypes *in vitro* as a function of age and time in culture, which may also foreshadow differences in outcomes when seeded in skin grafts and models for use *in vivo*.

## ACKNOWLEDGEMENTS

The authors thank the Comparative Medicine Institute, the NC State University and University of North Carolina at Chapel Hill Joint Department of Biomedical Engineering, and the NC State College of Veterinary Medicine for making this research experience possible.

## AUTHORS’ CONTRIBUTIONS

A.H.D. wrote the article, planned and executed the studies presented. K.M.P. and L.S.G. wrote the article, planned experiments, and assisted with execution of the studies presented. D.O.F and J.A.P. assisted in writing the article, planned experiments, and oversaw execution of the studies presented.

## SUPPLEMENTAL MATERIALS

**Supplemental Table 1.**
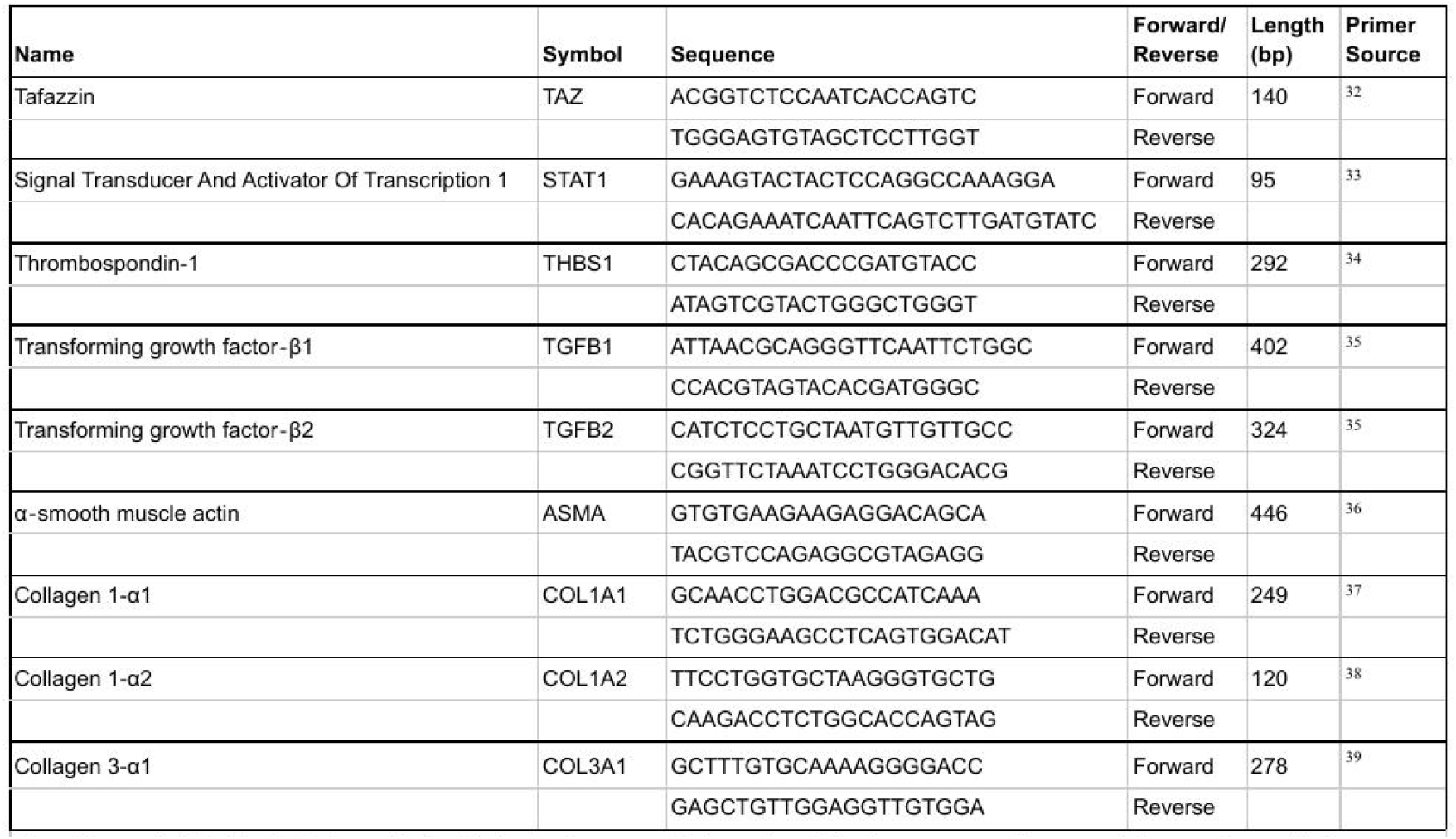
Primer Data. Selected genes of interest and their corresponding symbol are shown. Primer sequences, product length in basepairs (bp), and source are displayed as well.

**Supplemental Figure 1.**
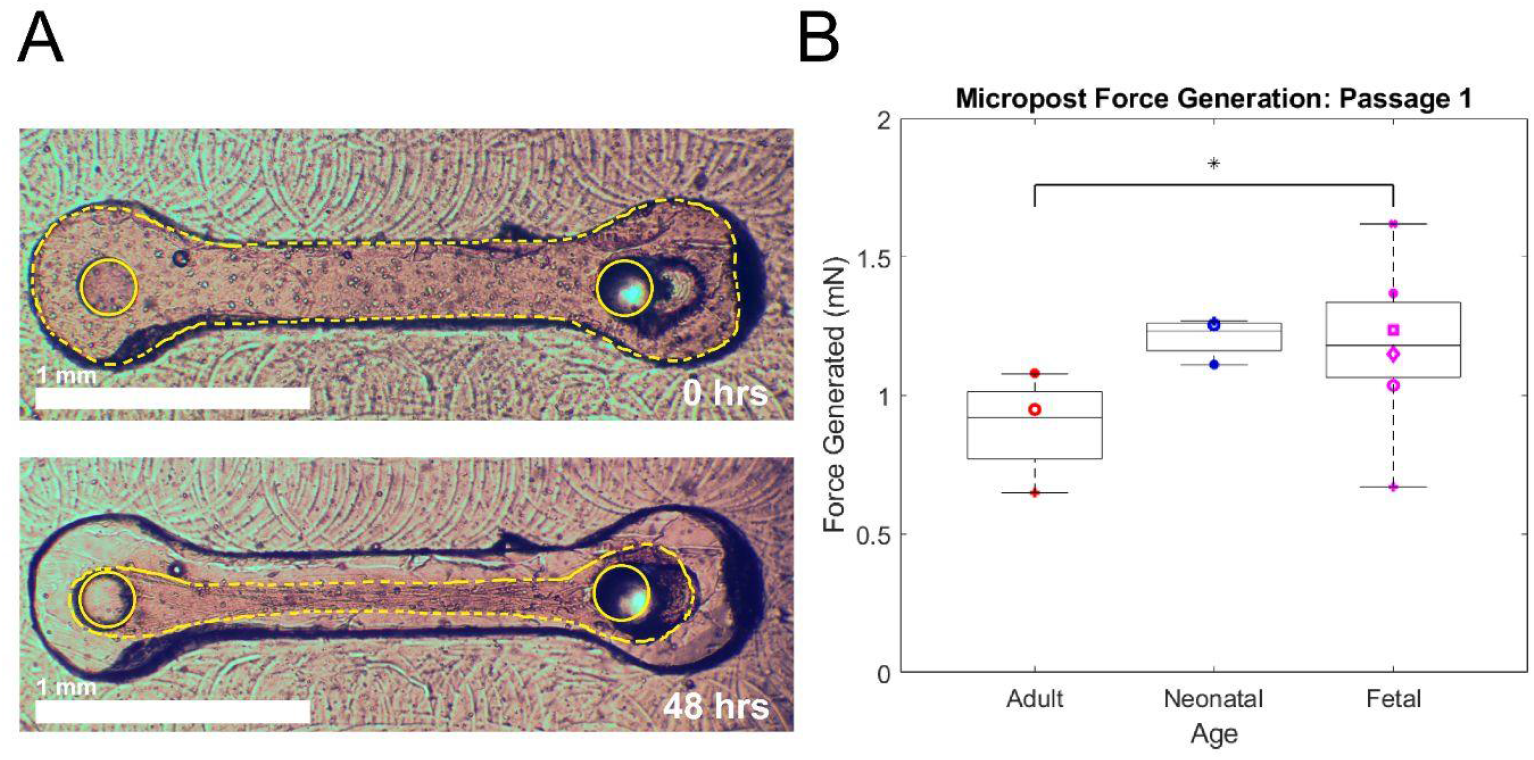
Fetal and neonatal age groups generated more force than adult in a micropost assay. (A) Fetal dermal fibroblasts suspended in a collagen hydrogel in the well of a PDMS micropost assay. Photos shown were taken immediately after seeding and 48 hours after seeding. The dotted yellow outline indicates the hydrogel, with white circles drawn on the top of each post. (B) Average micropost force generation in milli-Newtons for each cell sample in adult, neonatal, and fetal age groups. Adult and fetal force generation was significantly different when data from each micropost was analyzed in a two-tailed, unpaired t-test with an alpha value of 0.05. Cells from at least two different pigs with three technical replicates were analyzed in each group.

**Supplemental Figure 2.**
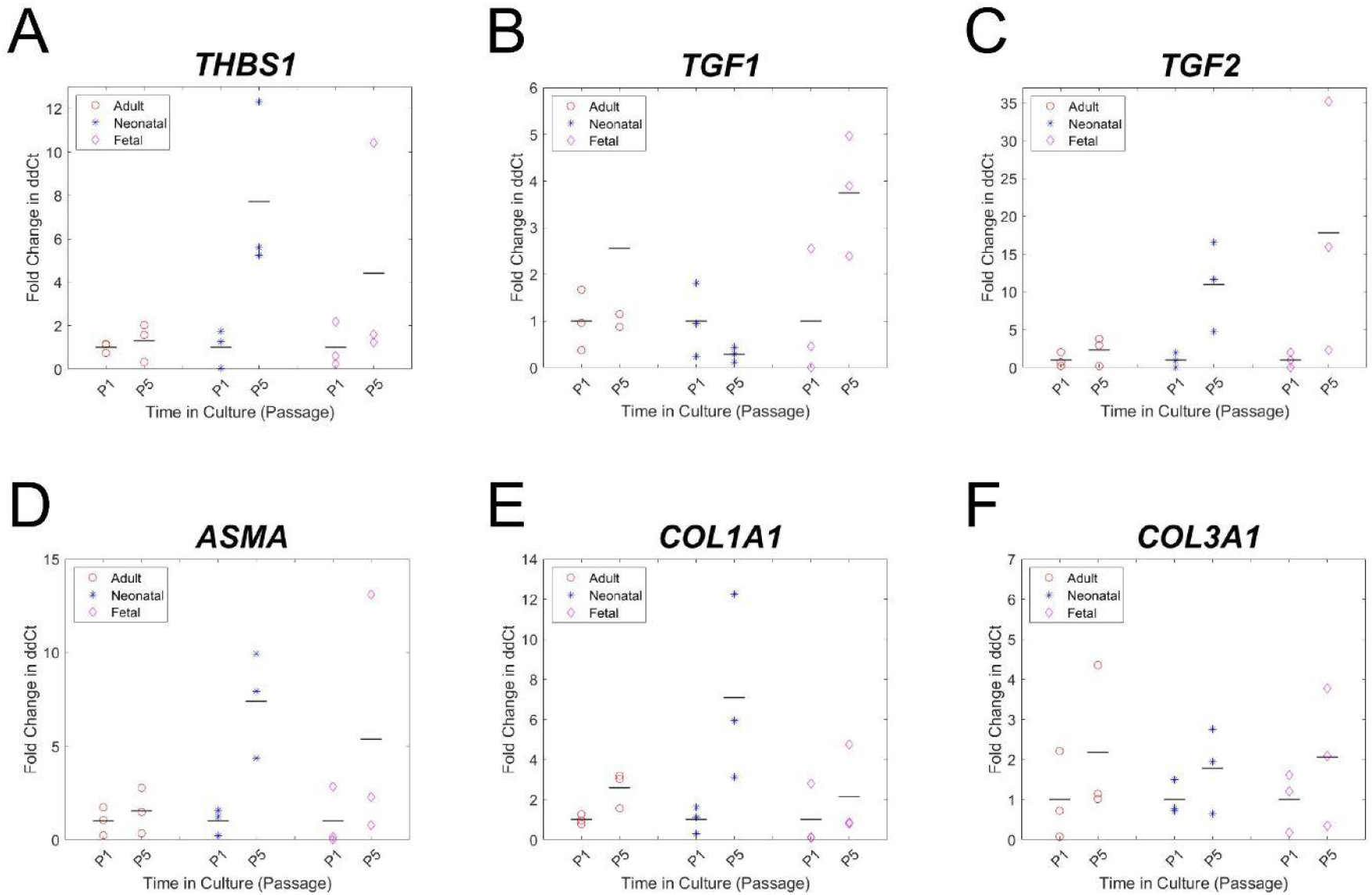
Gene expression analysis for a panel of genes related to wound healing, contraction, and ECM deposition. (A-F) Fold change in ddCt from passage 1 to 5 for adult (red circle), neonatal (blue cross), and fetal (magenta diamond) dermal fibroblasts. The mean for each group is indicated by a black bar. dCt values were standardized to GAPDH, followed by ddCt values where adult passage 5 was standardized to adult at passage 1, neonatal passage 5 to neonatal passage 1, etc. Cells from at least two different pigs with three technical replicates were analyzed in each group. No statistically significant differences were found in the genes in this figure.

